# Molecular Dynamics of DNA Translocation by FtsK

**DOI:** 10.1101/2022.01.03.474797

**Authors:** Joshua Pajak, Gaurav Arya

## Abstract

The bacterial FtsK motor harvests energy from ATP to translocate double-stranded DNA during cell division. Here, we probe the molecular mechanisms underlying coordinated DNA translocation in FtsK by performing long timescale simulations of its hexameric assembly and individual subunits. From these simulations we predict signaling pathways that connect the ATPase active site to DNA-gripping residues, which allows the motor to coordinate its translocation activity with its ATPase activity. Additionally, we utilize well-tempered metadynamics simulations to compute free-energy landscapes that elucidate the extended-to-compact transition involved in force generation. We show that nucleotide binding promotes a compact conformation of a motor subunit, whereas the apo subunit is flexible. Together, our results support a mechanism whereby each ATP-bound subunit of the motor conforms to the helical pitch of DNA, and ATP hydrolysis/product release causes a subunit to lose grip of DNA. By ordinally engaging and disengaging with DNA, the FtsK motor unidirectionally translocates DNA.

---

Many vital biological tasks such as viral DNA packaging, protein degradation, and chromosome segregation are completed by ring ATPase motors, which motors harness the energy from ATP binding, hydrolysis, and product release to translocate their biopolymer substrate through the central pore of the motor [1–6]. These motors have evolved mechanisms to coordinate activities within and between individual subunits, so that the assembly functions as a cohesive unit rather than a collection of subunits operating independently [7–9]. These mechanisms help all aspects of motor function. For instance, the motor must ensure that ATP is not prematurely hydrolyzed, that subunits are tightly gripping substrate before applying force, and that the resetting subunits are not tightly gripping substrate as they move in the direction opposite translocation. Although each aspect is likely controlled by distinct mechanisms, the motor nevertheless ensures that they occur in the proper sequence to yield unidirectional translocation of substrate. This implies that the motor has found ways to temporally connect individual mechanisms into one overall “well-oiled” mechanism.

Understanding how ATPase motors transition between different structural states within this integrated mechanism has been a subject of intense investigation. Because motor translocation is a dynamic process, techniques like single-molecule force spectroscopy have been particularly useful to characterize intermediate states and the kinetics of transition [10–12]. However, it is difficult to infer the underlying molecular mechanisms that produce force or promote coordination from singlemolecule experiments alone. To this end, researchers have utilized cryogenic electron microscopy (cryo-EM) to obtain atomic structures of several ATPase assemblies actively translocating their substrates, which in most cases cannot be accessed by crystallographic methods. Recently obtained structures include the bacteriophage φ29 DNA packaging motor [13], katanin [14], 26S proteosome [15], Lon protease [16], Vps4 [17], and FtsK [18]. Despite varying topologies and evolutionary divergences, each structure shows that the ATPase domains of actively translocating assemblies adopt helical arrangements as they engage with the biopolymer substrate. This strong conservation of quaternary structural arrangement, in addition to conservation of functional motifs (such as arginine fingers [8] and Walker motifs [19]), implies that there may be underlying mechanisms common to all translocating ring ATPase assemblies.

Among the aforementioned systems, bacterial FtsK is unique as it is the fastest DNA translocase currently known [20,21]. As each additional layer of control incurs additional activation energies that slow processes down, the mechanisms that regulate and coordinate translocation in FtsK may represent the minimal required mechanisms that are built upon by the broader class of ring ATPases. The FtsK translocase machinery is a homo-hexameric ring of ATPase subunits, where each subunit is further organized into domains. The α- and β-domains form the main translocase motor (called FtsK_αβ_), where the β-domains are the ATPase domains and DNA is translocated in the direction from the β-ring towards the α-ring.

Recently, Jean et al. [18] solved the cryo-EM structure of FtsK_αβ_ actively translocating double-stranded DNA, and showed that the motor is half occupied with the ATP analog and half occupied with ADP. ATP-analog-bound subunits were found to donate residues from “loop I” and “loop II” within the β-domain to grip opposite strands of DNA, whereas ADP-bound subunits do not. From this structure and additional biochemical evidence, they proposed a translocation model in which a hydrolysis event within a single subunit triggers conformational changes in every subunit. In this model, hydrolysis causes a subunit to disengage from DNA, permitting the other two upstream ATP-bound subunits to compact their α-/β-domains, thereby translocating DNA. At the same time, ADP release and ATP binding in a further upstream subunit causes that subunit to tightly grip DNA. These hydrolysis and nucleotide exchange events occur ordinally around the ring to yield continuous, step-wise translocation of DNA. While the domain-level movements are strongly supported by the structural data and agree with previous characterization of this system [22–24], the static structure offered little insight into the molecular interactions that give rise to such a mechanism. In particular, the nature of the driving force of subunit compaction during translocation, how subunits lose and gain registry with DNA based on nucleotide occupancy, and what mechanisms promote ordinal nucleotide exchange remain unknown.

Here, we used all-atom molecular dynamics (MD) simulations to investigate these open questions on molecular mechanisms governing DNA translocation by FtsK_αβ_. We advance the previously proposed model by predicting specific residue-level mechanisms that couple DNA gripping to ATP binding, subunit compaction to nucleotide occupancy, and additional *trans*-acting residues that may help promote ordinal ATPase activity. While the overall mechanism differs from those proposed for other systems [25,26], we observe the conservation of common functional elements across a broad class of molecular motors.

## Results

### ATP binding engages DNA gripping by loop II of the β-domain

To investigate conformational changes induced by bound nucleotide that contribute to DNA translocation, we performed microsecond-long MD simulations of subunit monomers in the apo, ATP-, and ADP-bound states. By understanding how an individual subunit responds to nucleotide occupancy, we can better understand how an assembly of subunits may respond to binding, hydrolysis, and product release at a single subunit.

The simulations revealed two distinct conformations of the ATPase active site depending on the state of the bound nucleotide. In the ATP-bound simulations, the catalytic glutamate (Glu596) points its carboxylate group towards the bound nucleotide, poised for catalytic activity (**Fig. 1A**). However, in the ADP-bound (as well as apo) simulations, Glu596’s carboxylate group instead hydrogen bonds with the phosphosensor motif (Thr630 and Gln631) (**Fig. 1B**). A result of this interaction is a decreased distance between the backbone of the catalytic glutamate and phosphosensor motifs, bringing DNA-gripping loop II Arg632 closer to Asp599 (**Fig. 1C**). In the ATP-bound simulations, Arg632 points away from the surface of the protein, such that it can potentially engage with substrate DNA (**Fig. 1A**); in the apo and ADP-bound simulations Arg632 is flush against the surface of the protein, interacting with Asp599, and would therefore not be expected to engage with DNA (**Fig. 1B**).

**Figure 1.**
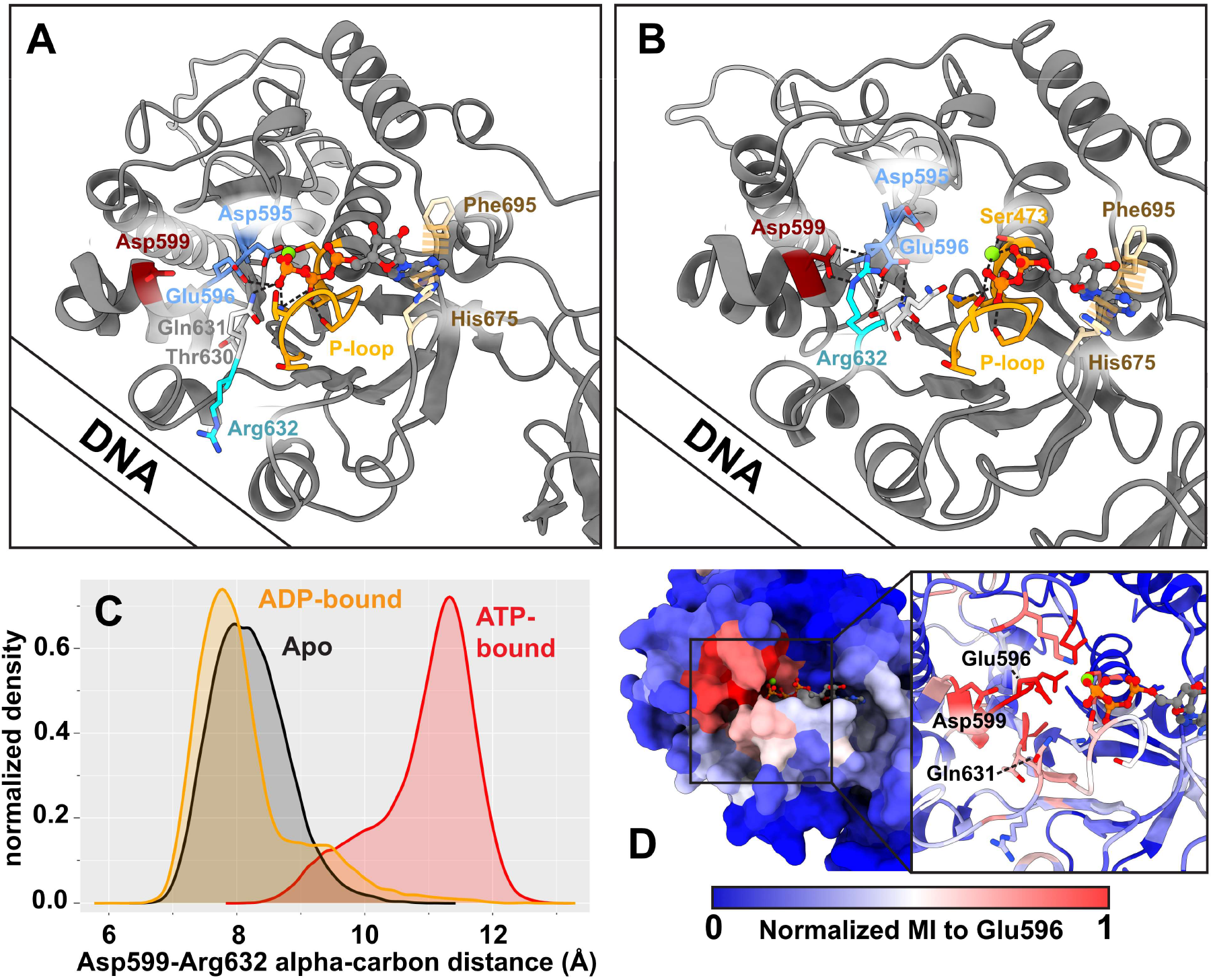
ATP-binding causes loop II to grip DNA. **(A)** ATP-bound conformation predicted from MD simulations. The Walker A motif (P-loop) is orange, Walker B motif is blue, the sensor motif is white, DNA gripping loop II Arg632 is cyan, Asp599 is dark red, and residues that π-stack with adenosine are tan. **(B)** ADP-bound conformation predicted from MD simulations. Arg632 is held by Asp599 and cannot grip DNA. The approximate position of DNA is indicated. **(C)** In absence of the γ-phosphate of ATP, catalytic Glu596 interacts with the sensor motif (Thr630-Gln631), bringing Arg632 closer to Asp599. **(D)** Mutual information cluster extracted from MD simulations. The catalytic Glu596 has high MI with residues predicted to coordinate the γ-phosphate, and the MI decays across the Walker A motif binding pocket. Thus, this coordination primarily senses the γ-phosphate and alters the conformation of Arg632 to either grip DNA or ungrip DNA.

The dependence of the catalytic glutamate on the presence/absence of the γ-phosphate of ATP to control DNA-gripping is reminiscent of the “glutamate switch” found in AAA+ enzymes [9,27]. We previously predicted *via* mutual information extracted from MD simulations that viral DNA packaging enzymes also use a glutamate switch to couple ATPase and DNA packaging activities, a prediction that was corroborated by functional mutagenesis [28]. Motivated by these results, we extracted mutual information from our FtsK simulation trajectories using the CARDS (<u>C</u>orrelation of <u>A</u>ll <u>R</u>otameric and <u>D</u>ynamical <u>S</u>tates) method [29] to see if a similar connection is predicted in FtsK. Our simulations revealed that the catalytic glutamate residue has high mutual information with several residues that are expected to coordinate the γ-phosphate of ATP (**Fig. 1D**). Additionally, this mutual information extends to Asp599, Gln631, and Thr630, which can control the position of Arg632. This supports the hypothesis that nucleotide-occupancy is sensed by the catalytic residues and relayed to loop II DNA-gripping residues, namely Arg632. In sum, our simulations predict a feasible mechanism for how FtsK regulates its grip on DNA pre- and post-hydrolysis that is consistent with other translocating ATPases.

### Rotations of the α- and β-domains are the primary modes of motion

We calculated the principal components of motion from our monomer simulations and found that the dominant motion is a rocking of the β-domain relative to the α-domain (**Fig. 2A**). This is perhaps expected, as the two domains are connected by a flexible linker and this motion was proposed to be responsible for translocating DNA based on structural evidence [18]. However, while monomer dynamics can provide insights into the predominant motions of a subunit, these motions can be altered by interactions with neighboring subunits and DNA. Thus, in addition to our monomer simulations we also performed long timescale MD simulations of the hexameric assembly with DNA in the central pore. PCA of the entire hexameric complex predicted that subunit motions are coordinated through their association with DNA. The four subunits that grip DNA rock their domains sympathetically, whereas the two subunits not engaged with DNA do not exhibit such correlation (**Fig. 2B, Movies S1 and S2**). Thus, maintaining helical registry with DNA allows subunits to cooperate in order to translocate DNA.

**Figure 2.**
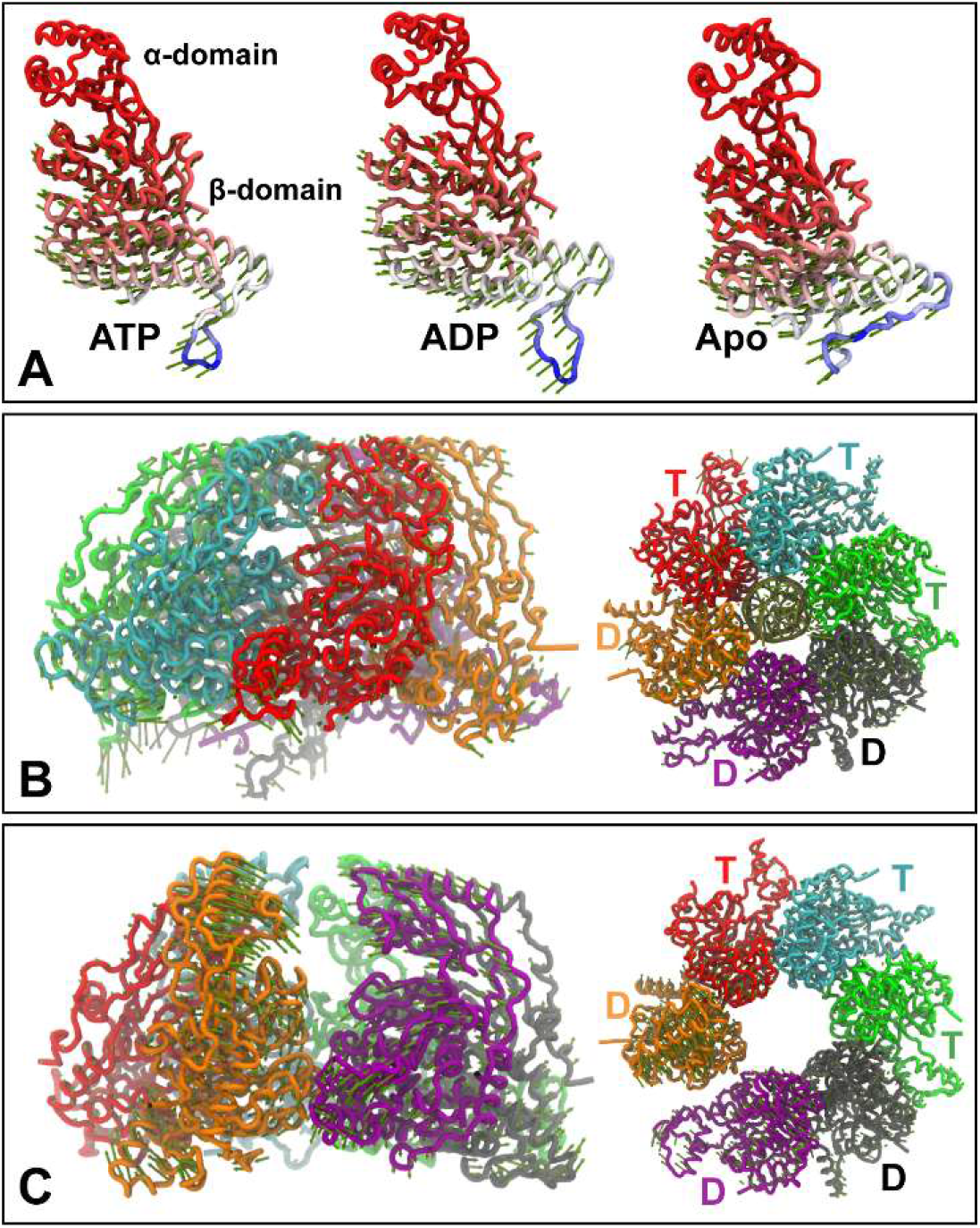
Principal components of motion. **(A)** The first principal component of motion predicted from monomer simulations in the ATP-, ADP-bound, and apo states after aligning to the α-domain is a rocking of the β-domain. This motion has been proposed to be the translocation mechanism. Proteins are colored by root-meansquare fluctuation (RMSF), blue resides have high RMSF, red residues have low RMSF. The first principal component predicted in the hexamer simulation with DNA. The sympathetic motions are more clearly observed in Supplemental Movies S1 & S2. Subunits are labeled with a “T” if they are ATP-bound, and “D” if they are ADP-bound. **(C)** The first principal component predicted in the hexamer simulation without DNA.

To further investigate if substrate DNA is necessary to promote coordinated motions of subunits, we performed MD simulations of the hexamer assembly with no DNA in the pore. These simulations predicted that the ring begins to open at the interface between the ADP-bound DNA tracking subunit (orange subunit in **Fig. 2C**) and a non-DNA-tracking subunit (purple subunit in **Fig. 2C**), suggesting that interaction with DNA is necessary to stabilize the quaternary structure of the ATPase assembly, in agreement with experimental data [24]. Thus, it is possible that DNA also plays a regulatory role in its own translocation by providing stabilizing interactions for the ATPase ring.Next, we investigated the principal components of motion of individual subunits within the hexamer from the DNA-bound simulations. Our analysis predicted that, like their monomeric counterparts, dynamics of a monomer within the hexamer are also dominated by rocking of the α- and β-domains. However, the first principal component of motion also encompasses a shear between the α- and β-domains. Based on this observation, we defined two collective variable angles: *extension*, the angle that describes the separation between DNA-gripping motifs in the α- and β-domains, and *twist*, the angle that describes the shearing between the domains (**Fig. 3A**). We then projected our hexamer simulations onto these two collective variables and found that there is a continuous distribution of states sampled by the hexamer (**Fig. 3B**). The subunits with the largest extension and twist are those bound to ATP, which track the helical DNA substrate. The subunits with less extension and twist are ADP bound, which are largely not constrained by interaction with substrate DNA. This suggests that the extended states are likely stabilized by interaction with DNA.

**Figure 3.**
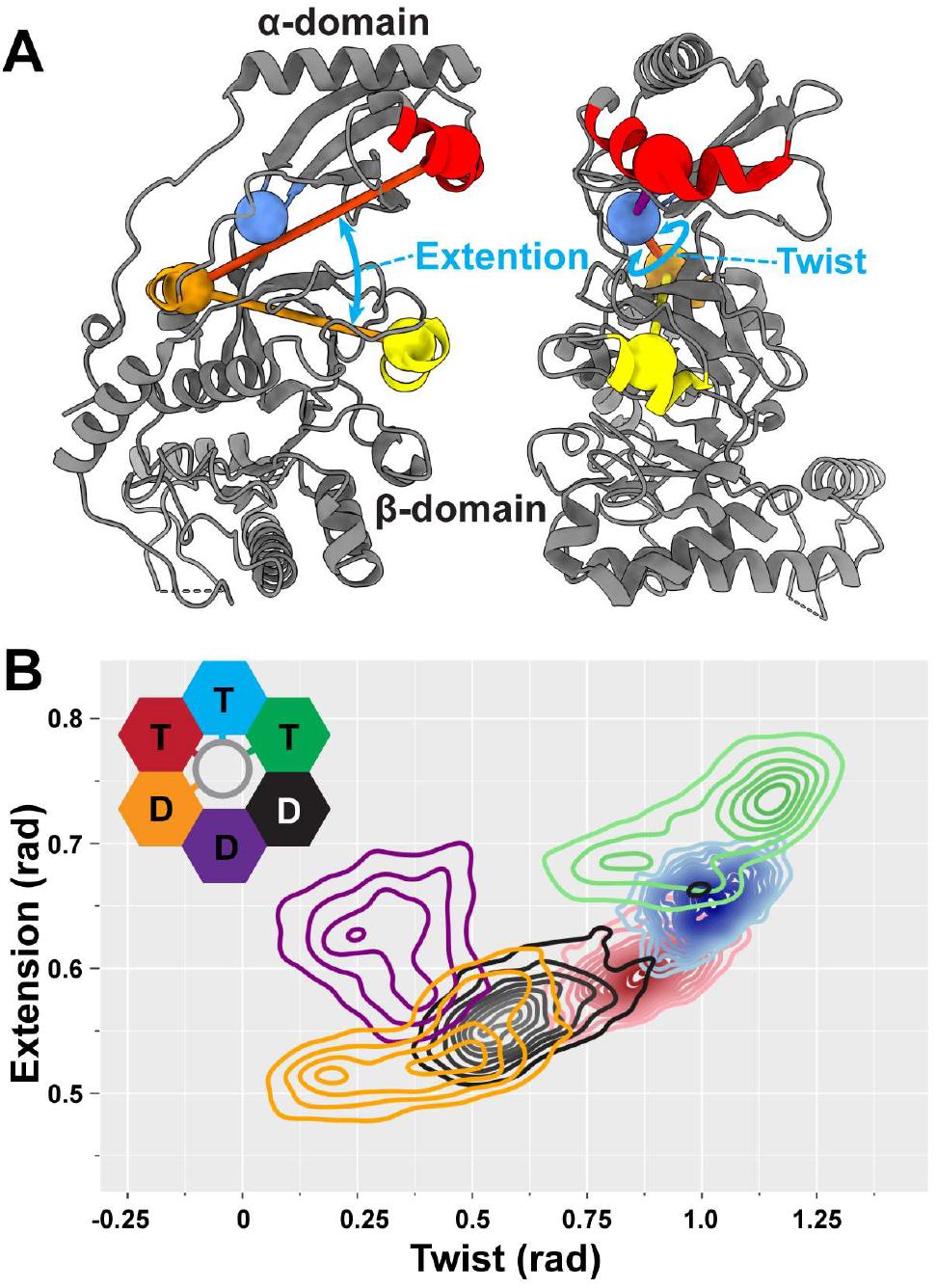
Collective variables that characterize motion. **(A)** The extension angle describes the separation of DNA-gripping helices in the α- and β-domains. The twist angle describes the shearing between these helices. **(B)** Subunit conformations sampled in the hexamer simulation with DNA projected on the twist and extension angles. DNA-tracking subunits (green, cyan, red, orange) form a continuous distribution. Subunits that do not grip DNA (purple, black) do not necessarily follow this trend, as they are not constrained by interaction with DNA.

### Nucleotide binding promotes compaction

To further probe the connection between nucleotide occupancy and preferred relative orientations of the α- and β-domains, we calculated free-energy landscapes of a subunit in the extension and twist collective variable space using well-tempered metadynamics simulations. Our calculations predict that both ATP- and ADP-bound subunits prefer a compact conformation (**Fig. 4A**). On the other hand, the apo state subunit samples relatively broad collective variable space with little energetic penalty (**Fig. 4A**). In other words, the nucleotide-bound states are predicted to be rigid and compact, whereas the apo state is predicted to be flexible.

**Figure 4.**
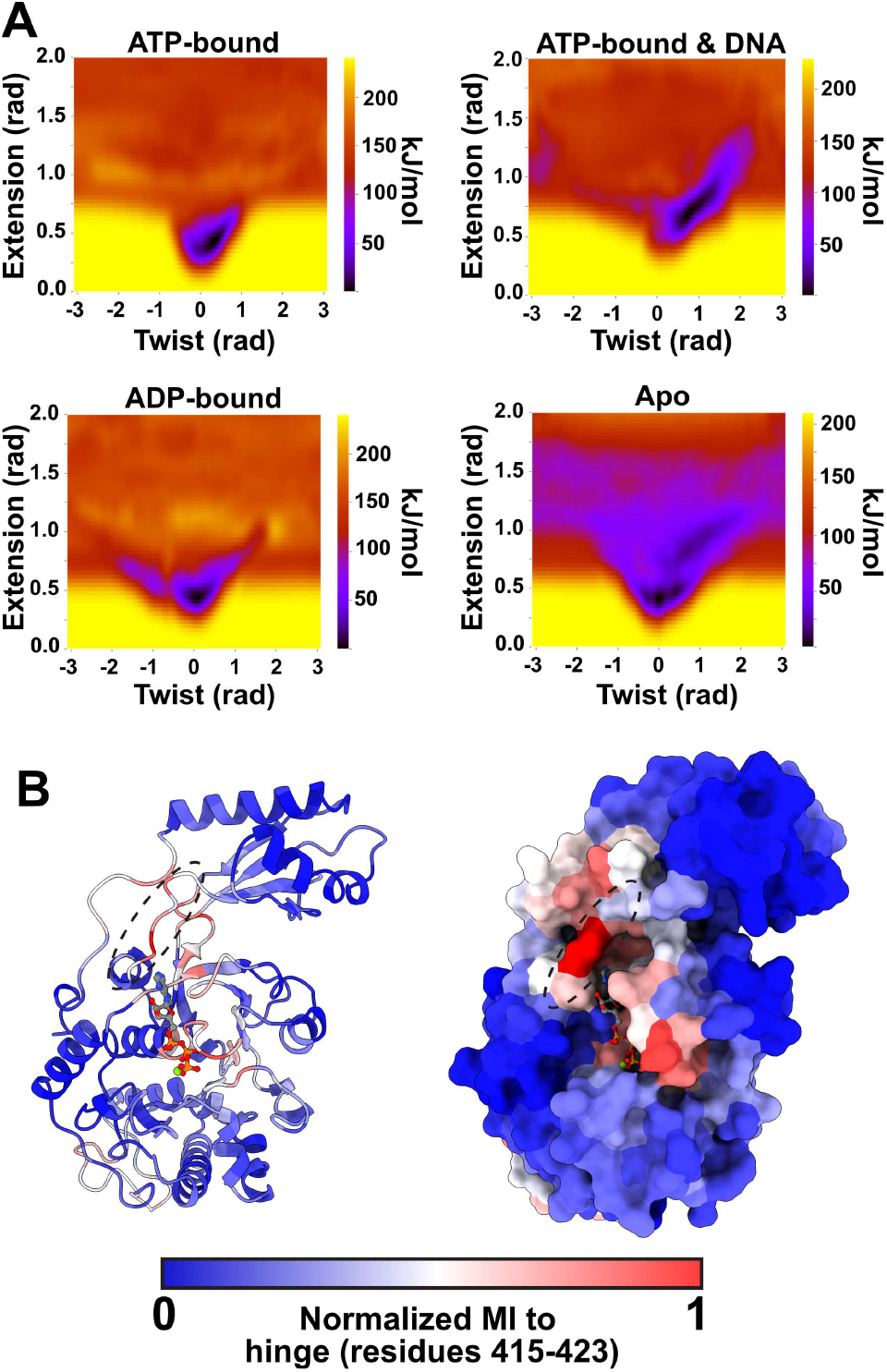
Free-energy and mutual information calculations predict that nucleotide compacts subunits. **(A)** Free-energy landscapes in the twist-extension collective variable space calculated by well-tempered metadynamics. In the nucleotide-bound states, the compact state (low extension) is the global free-energy minimum. In the apo state, while the compact state is the global minimum, the collective variable space is much more accessible, with metastabilities emerging in twisted and extended conformations. DNA stabilizes twisted and extended conformations in the ATP-bound state as the subunit tracks the helical phosphate backbone. The calculated free-energy minimum corresponds well with the states sampled in by DNA-gripping subunits in the hexamer simulation (Fig. 3B). **(B)** Mutual information calculated by CARDS, targeting the hinge connecting the α-/β-domains. Hinge (dashed circle) and interface residues have high mutual information to residues that bind the adenosine and α-/β-phosphates of ATP. Notably, this mutual information decays prior to reaching the Walker B motif, which primarily responds to the γ-phosphate of ATP, suggesting that compaction is a hydrolysis-independent process.

Next, we wondered how interaction of a subunit with DNA could affect this free-energy landscape. Thus, we likewise calculated the free-energy landscape of an ATP-bound subunit complexed with DNA. We found that interaction with DNA stabilizes the more twisted and extended conformations (**Fig. 4A**). The shape and span of this free-energy minima valley agrees well with the states sampled by ATP-bound monomers in our hexamer simulation (**Fig. 3B**). Thus, the interaction energy gained from protein-DNA interactions could stabilize ATP-bound subunits in twisted and extended conformations.

Lastly, to determine how nucleotide occupancy might control the conformation of the subunit, we extracted mutual information from our equilibrium MD simulations. We performed a target site analysis relative to the flexible linker (or hinge) connecting the α-domain to the β-domain, reasoning that controlling the linker between the domains could provide a means to control the relative orientations of the two domains. We find that this hinge has high mutual information to residues involved in binding the adenosine ring (e.g., His675 and Phe695) and the α-/β-phosphates (e.g., the Walker A motif/P-loop) (**Fig. 4B**). Notably, there is little mutual information shared with the Walker B motif, which primarily responds to the presence or absence of the γ-phosphate of ATP. Thus, the shared information is coordinated by ATP and ADP alike, explaining why both nucleotides drive compaction.

### Geometry of the β-ring promotes ordinal activity

Based on the solved cryo-EM reconstruction of actively translocating FtsK_αβ_, the most commonly observed state of the motor is contiguous stretches of three ATP-bound and three ADP-bound subunits. Because the class-averaged structure is dominated by this one configuration despite potentially stalling at any translocation step, it was inferred that each translocation step results in a rotationally symmetric configuration [18]. Additionally, during translocation, subunits must release ADP to bind ATP. This implies that the motor has evolved mechanisms that selectively promote ordinal ADP release (and subsequent exchange with ATP) in one subunit and not the other two ADP-bound subunits, otherwise different ATP/ADP nucleotide permutations would have been observed.

Our MD simulations of the hexameric assembly with substrate DNA sheds light on one possible mechanism. At the proposed nucleotide exchanging interface (and not in the other ADP-bound interfaces) additional transacting residues interact with the bound ADP (**Fig. 5A,** green subunit donating trans-acting residues into the black subunit). Specifically, we find that Arg548 interacts with the phosphates of ADP, and Asn549 with the adenosine base (**Fig. 5B**). These interactions might destabilize the interactions between ADP and cis-acting residues, thus potentially promoting ADP release and subsequent ATP binding.

**Figure 5.**
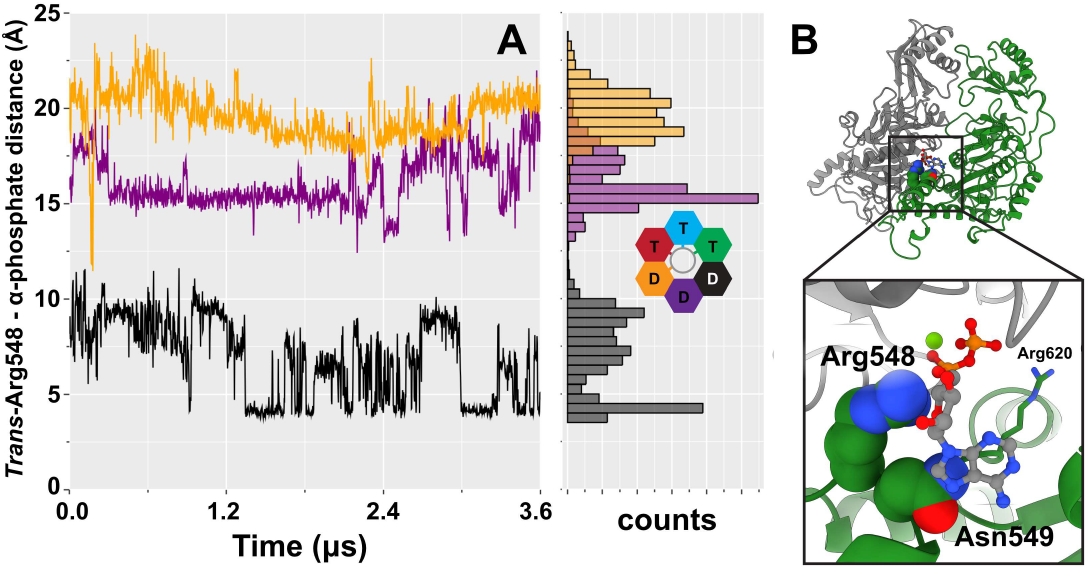
Geometry of the ring promotes ordinal ADP release. **(A)** Simulated distance between the guanidine carbon of Arg548 and the α-phosphate of ADP in the neighboring subunit. Arg548 only interacts closely with the α-phosphate of the subunit next in line to exchange ADP for ATP, suggesting that it helps promote nucleotide exchange. Traces are colored according to the *cis* binding site. **(B)** A depiction of this interaction. The canonical *trans-act*ing arginine finger, Arg620, is also shown.

The interactions between Arg548/Asn549 and the neighboring subunit’s ADP seem to be enforced by the geometry of the ring. The residues are donated by the most extended ATP-bound subunit. Because subunit extension is not a simple linear translation, but rather is accomplished by rocking the β-domain relative to the α-domain, there is a secondary effect of extension pushing trans-acting residues upwards and towards the neighboring subunit (**Fig. 2 and Movie S3**). Thus, a subunit binding ATP and engaging with DNA at the bottom of the β-ring helix positions Arg548 to act as an exchange residue, promoting ADP release and ordinal nucleotide exchange.

A similar mechanism could potentially ensure ordinal ATP hydrolysis events. In the solved cryo-EM reconstruction, the canonical arginine finger (Arg620) is near to the γ-phosphate of ATP in every ATP-analog-bound subunit. Hence, it is not readily apparent why hydrolysis would only occur in one specific subunit and not the other two ATP-bound subunits. In our hexamer simulation, Arg548 also transiently interacts with the γ-phosphate of ATP in addition to the canonical arginine finger interaction (**Fig. S1**). Although this interaction is less stable than the ADP-bound interaction shown in Figure 5 and occurs in an unexpected subunit, it nevertheless demonstrates the possibility for this residue to contribute towards hydrolysis. In support of this, our PCA results show that this donation is the primary mode of motion at the hydrolyzing interface when aligned about the α-domain (**Movie S3**). Thus, we propose that this second arginine residue may also play a catalytic role in addition to the canonical arginine finger to help ensure ordinal ATP hydrolysis activity.

### Distortion of the α-ring may help DNA translocation

Whereas the β-ring forms a partial helix as subunits track substrate DNA, the α-ring is planar and thus cannot track substrate DNA. In the cryo-EM reconstruction, the α-ring is essentially a symmetric hexamer (**Fig. 6A**). This arrangement is similar to viral DNA packaging ATPases, where the ATPase domain forms a helix as they track DNA, but the C-terminal domain remains a symmetric planar ring connected to the portal assembly [13]. This planar ring had been previously proposed to act as an aperture that guides the positioning of DNA through the pore and promotes high processivity in the FtsK_αβ_ assembly [18].

**Figure 6.**
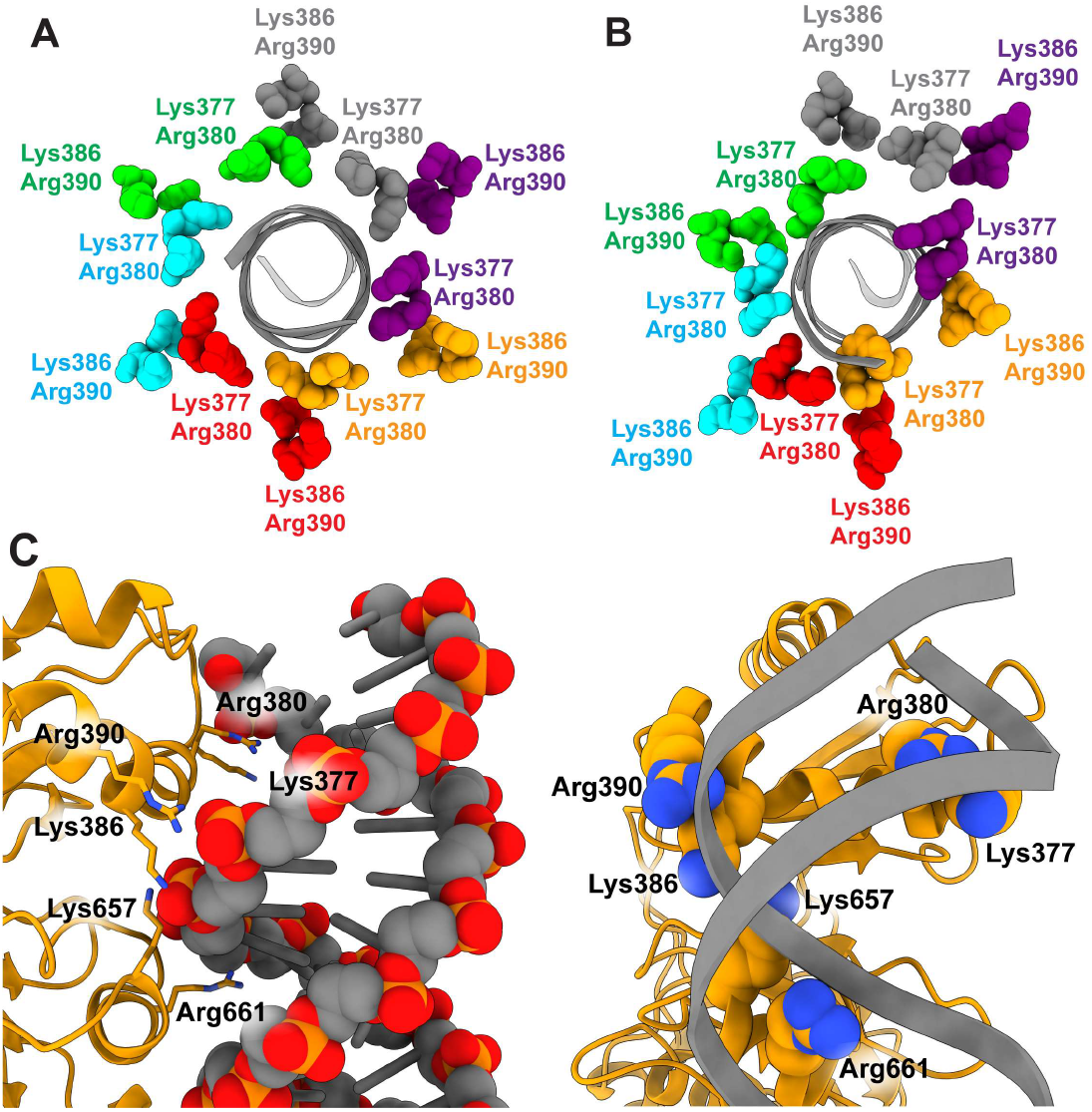
The α-ring can deform to promote DNA-binding. **(A)** The cryo-EM reconstruction (PDB: 6T8B) shows that the α-ring is symmetric around substrate DNA. Two α-domain motifs are shown as spheres, Lys377/Arg380, which interacts with DNA, and Lys386/Arg390, which does not. **(B)** MD simulations predict that the α-ring can distort, such that the ADP-bound subunit at the top of the β-ring helix can grip DNA with both α-domain motifs. These interactions are shown in **(C)**. The geometry of the α-domain allows the subunit to track both strands of DNA. The new interactions of Lys386/Arg390 may compete with the grip of loop I Lys657 in the β-domain, allowing the subunit to lose grip of DNA in the β-domain before resetting to the bottom of the helix.

In our hexameric simulations, we find that the α-ring can distort to promote more DNA gripping residues to contact DNA (**Fig. 6B**). Notably, the compact hydrolyzing (orange) subunit twists to promote new contacts with the DNA phosphates through Lys386 and Arg390, in addition to the contacts made by Lys377 and Arg380 observed in the cryo-EM reconstruction (**Fig. 6C**). The geometry of the DNA-gripping motif of the α-domain is such that Lys377/Arg380 grip one strand of DNA, while Lys386/Arg390 grip the other, similar to how loop I and loop II in the β-domain grip opposite strands. Because this subunit is the most compact, these additional contacts compete for phosphate grip with the loop I DNA-gripping lysine (Lys657) (**Fig. 6C**). Thus, as hydrolysis causes this subunit to lose registry with DNA in loop II (**Fig. 1**), compaction of this subunit can exchange its DNA grip between loop I and the α-domain.

We note that the transient behavior of the interactions between Lys386/Arg390 and DNA in the simulations, and the fact these interactions are not observed in the cryo-EM reconstruction, suggest that the interactions may represent a short-lived state that helps the β-domain completely lose grip of DNA before resetting back down the helix. In this way, handing off grip of DNA from the β-domain to the α-domain coupled to ATPase activity and subunit compaction is a plausible mechanism by which the motor translocates DNA.

## Discussion

### Proposed FtsK mechanisms

The work presented herein expands upon the model proposed in Jean et al. [18]. In the proposed model, simultaneous conformational changes occur around the ring coupled to ATP hydrolysis at the “top” of the β-ring and nucleotide exchange at the “bottom” of the β-ring (**Fig. 7,** orange and black subunits). These conformational changes cycle the overall assembly configuration one subunit around the ring. However, several fundamental questions remain about the molecular nature of these events. Our results allow us to address these questions by proposing specific molecular mechanisms that coordinate activities across subunits and generate force necessary to translocate DNA.

**Figure 7.**
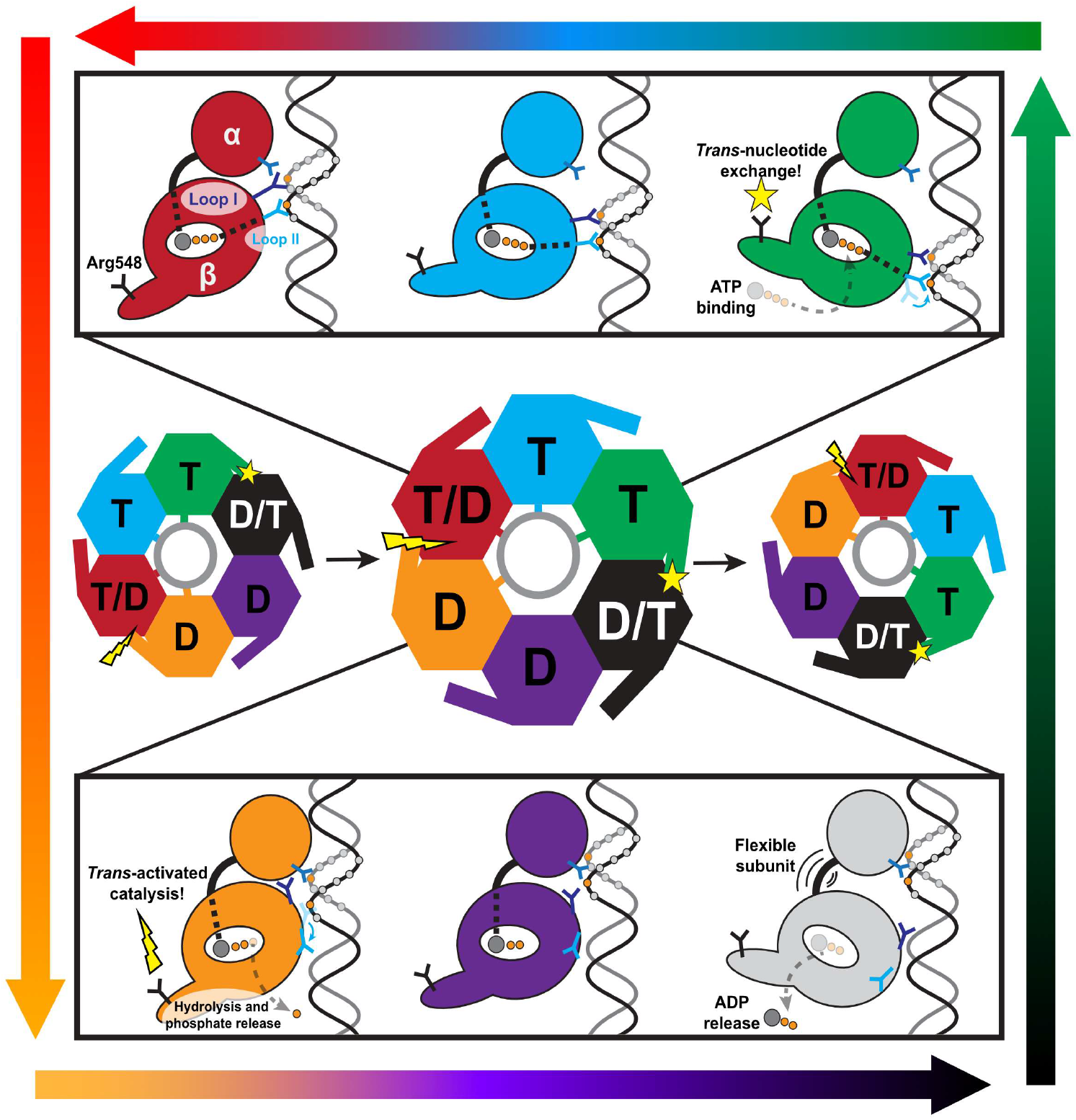
Overall mechanochemical cycle of DNA translocation by FtsK. The overall mechanochemical model is based off the one proposed in Jean et al. [18]. Here, dynamic information taken from MD simulations is used to flesh out the individual steps in the cycle. In the center, the conformational changes of the hexameric complex are depicted. Subunits are labeled “T” if they are ATP-bound, “D” if they are ADP-bound, “T/D” if that subunit is expected to hydrolyze ATP to ADP, and “D/T” if that subunit is expected to exchange ADP for ATP. Trans-catalyzed ATP hydrolysis and nucleotide exchange are shown by a lightning bolt and star, respectively. Contacts with DNA (gray circle) are shown as sticks. In the top panel, the ATP-bound subunits are depicted. Signals sent from the bound nucleotide to Loop I and the flexible hinge are shown as dashed lines. For simplicity, the multiple DNA contacts in Loop I, Loop II, and the α-domain are shown as singular prongs. In the bottom panel, ADP-bound subunits are shown, and the “D/T” subunit is shown as passing through the transient apo state. In the top and bottom panels, some DNA phosphates are shown as circles colored orange if they were in registry with the subunit and gray if they were not. These phosphates guide the eye to DNA translocation as a single subunit begins at the bottom of the ATP helix (green); works its way towards the top while remaining in registry with and translocating DNA (cyan, red); passes grip from the β- to the α-domain in the post-hydrolysis ADP-bound state (orange); and resets to the bottom of the β-domain helix while the subunit is not engaged with DNA (purple and gray), finally realigning with the phosphate backbone and beginning the cycle over again (green). This order of events is guided by the gradient arrows encircling the figure.

We begin our description of the model at the configuration solved in Jean et al. [18]. In this configuration, three ATP-bound subunits’ β-domains track DNA substrate. Our simulations show that while bound nucleotide drives subunits into a compact conformation, the extended subunits at the bottom of the β-ring twist and are stabilized by interactions with the double-stranded DNA substrate (**Fig. 4**). At the top of the β-ring helix sits the hydrolyzing subunit; ATP hydrolysis signals DNA-gripping loop II to disengage with DNA (**Fig. 1, Fig. 7**). Because this subunit is the most compact, DNA-gripping loop I comes into direct contact with the α-ring, passing DNA grip from the β-domain to the α-domain (**Fig. 6**). Thus, hydrolysis and compaction cause the subunit at the top of the β-ring helix to lose registry with substrate DNA in its β-domain.

At the other end (bottom) of the β-ring helix, ADP release is promoted *in trans* by an ATP-bound subunit (**Fig. 5, Fig. 7**). During the transient apo state, this subunit becomes highly flexible (**Fig. 4A**), allowing it to extend to the bottom of the β-ring helix, potentially promoted by nascent interactions with DNA phosphates. Subsequent ATP binding causes this subunit to engage tightly with DNA, stabilizing it at the bottom of the β-ring helix. The net effect of a subunit losing registry with DNA at the top of the helix and a subunit gaining registry with DNA at the bottom of the helix is a compaction of the remaining ATP-bound subunits, generating force to translocate DNA 2 base-pairs towards the α-ring.

### Overall conserved features of translocation

The fundamental elements of the proposed mechanism are:

1. Nucleotide binding drives two competing effects: strong interaction with DNA that forces subunits into a helical configuration, while simultaneously promoting compaction of a subunit through internal conformational changes. Helicity promoted by interaction with DNA wins the competition.
2. ATP hydrolysis causes a subunit to disengage with DNA, which allows neighboring subunits to undergo ATP-induced compaction, translocating DNA past the hydrolyzing subunit.

Despite different aesthetic appearances of the models, these key features are strikingly similar to the fundamental features in the helical-to-planar model proposed for viral DNA packaging ATPases [13,25,30]. Additionally, many AAA+ have been proposed to utilize a “handover-hand” model of substrate translocation [31,32]. In this model, ATP-bound subunits form a helical pitch as they track the biopolymer substrate. Hydrolysis at one end of the helix disengages a subunit from the substrate, and that substrate begins to reset to the other side of the helix. Translocation is accomplished by the remaining ATP-bound subunits shifting their positions as they maintain grip with the biopolymer substrate. Thus, it may be possible that these models are converging to the same fundamental aspects outlined above, and that differences emerge as system-specific constraints modulate the overall mechanism.

### Differentiating features of FtsK translocation and viral DNA packaging

While there may be mechanisms common to all translocating ring ATPases, each system has a unique evolutionary history and system-specific optimizations that must be considered. For this discussion, we will contrast the FtsK model to the viral DNA packaging helical-to-planar model.

The pore of the hexameric FtsK_αβ_ assembly is larger than the diameter of dsDNA, such that some subunits do not engage with DNA [18]. This allows for FtsK_αβ_ subunits to reset from one end of the helix to the other during translocation without fighting against the translocated DNA. Thus, the pore size permits a continuous translocation mechanism as subunits are free to both translocate and reset simultaneously. On the other hand, viral DNA packaging ATPase rings are pentameric assemblies when attached to procapsids and actively translocating DNA, likely maintaining symmetry with the unique fivefold vertex of the capsids at which they assemble [13,33,34]. The pore of the pentameric ATPase assembly is roughly the same diameter as dsDNA. Thus, any subunit resetting from one end of the helix to the other would have to fight against the translocated DNA. Accordingly, viral packaging ATPases have adapted burst-dwell packaging dynamics [11,30,35]; DNA is translocated one helical turn in during a quick burst that is followed by prolonged dwells wherein no DNA is translocated as the entire motor resets. Partitioning translocation and motor reset phases ensures that each subunit is moving in the same direction and subunits do not compete against each other. Based on these observations, we propose that pore-size and the observation of burst-dwell dynamics can be used as discriminating whether a motor assembly utilizes a FtsK-like or a helical-to-planar translocation mechanism.

A second difference is the topology of the linker motif. In AAA+ and viral packaging ATPases, the connector between the globular N-terminal (ATPase) domain and C-terminal domains is a α-helical-rich lid subdomain that mediates a large amount of inter-subunit interactions [14,25]. In the helical-to-planar and hand-over-hand models, rotation of the lid subdomain was proposed to be the force-generating mechanism [25,26]. On the other hand, the connector between the α-/β-domains of the FtsK assembly is a short coil and does not interact with neighboring subunits, and thus an analogous mechanism cannot be the force-generating step. This difference is perhaps related to the overall goals of viral packaging ATPases/AAA+ and FtsK-like motors. Viral packaging ATPases/AAA+ motors often must produce high forces to accomplish their tasks, such as packaging DNA to near crystalline density inside of a viral capsid [36], or disassembling microtubules [14]. A subunit being able to leverage neighboring subunits *via* the lid subdomain may help these motors generate high forces. On the other hand, FtsK-like motors do not translocate DNA against high forces but do translocate DNA an order of magnitude faster than viral packaging ATPases. Hence, FtsK-like motors may have adapted to favor mechanisms that promote fast translocation rather than those that generate high forces. Thus, the topology of the linker may be means to discriminate between mechanisms.

Lastly, hydrolysis is controlled by distinct means. In FtsK-like assemblies, hydrolysis is catalyzed *in trans* by two subunits that are helically offset from each other [18]; viral DNA packaging ATPases are proposed to catalyze hydrolysis upon a planar alignment of two subunits [13,25]. The difference in interfacial geometry at the time of ATP hydrolysis requires different sites for the canonical arginine finger within the subunits. Thus, the sequence location of the arginine finger may also be a means to discriminate between translocation models.

## Methods

### Hexamer simulations

Initial structures were taken from the atomic model built into the cryo-EM reconstruction of hexameric FtsK_αβ_ motor assembly captured in the middle of translocation, containing three ADP- and three ATP-γS-bound subunits (PDB: 6T8B) [18]. ATP-γS was manually changed to ATP. Missing loops (e.g., residues ~570-585) were rebuilt using MODELLER [37], using the high-resolution crystal structure of FtsK_αβ_ (PDB: 2IUU) as a constraint. Mg^2+^ was added to ATP/ADP binding sites where it was not built.

Equilibrium simulations of the hexameric complex were initialized in the AMBER18 simulation package prior to running on the Anton 2 supercomputer [38]. The protein interactions were described using the AMBER ff14SB force field [39], and DNA was described using the OL15 modifications [40]. ATP and ADP parameters were taken from the AMBER parameter database [41]. The hexamer and DNA assembly was centered in a cubic box of TIP3P water with 12 Å padding. Counter ions were added to make the system charge-neutral, and then additional salt was added to reach 150 mM concentration. The systems were subjected to 300 steps of steepest descent and conjugate gradient energy minimization. Subsequently, the systems were equilibrated using the GPU-accelerated pmemd module in AMBER18 [42] as follows. The SHAKE algorithm was used to constrain bonds connecting hydrogen atoms to heavy atoms, and a 2-fs integration timestep was used to propagate the equations of motion. The systems were heated slowly from 100 K to 310 K over the course of 100 ps in the canonical (NVT) ensemble. Then the systems were equilibrated in the isobaric-isothermal (NPT) ensemble at 310 K and 1 bar using the Monte Carlo barostat and Langevin thermostat for 50 ns. Finally, the hexamer system was simulated for 3.6 μs in the NPT ensemble on the Anton 2 supercomputer using the Nose-Hoover thermostat and MTK barostat, with a 2.5-fs integration timestep.

### Monomer simulations

Monomer simulations were started by isolating specific subunits from the hexameric complex: ATP-bound simulations started with an isolated ATP-bound subunit, and ADP-bound simulations started with an isolated ADP-bound subunit. The apo state simulations started from an ATP-bound subunit where Mg^2+^-ATP was manually removed from the binding pocket, hence, any differences predicted between ATP-bound and apo simulations are a direct result of nucleotide occupancy and not initial structures.

Because it has been shown that the TIP3P water model under-predicts protein-water interactions, and thus promotes compact states, we were concerned that monomer systems may be more susceptible to this bias without the constraints of neighboring subunits and DNA. We therefore chose to describe monomer systems with the recent AMBER ff19SB force field [43] with the OPC water model [44], which strengthens protein-water interactions and stabilizes extended states. The systems were centered in octahedral periodic boxes with OPC water molecules and 14 Å padding. The systems were energy-minimized and equilibrated as described above. Duplicate simulations were performed of each system diverging at the heating stage of equilibration. Production simulations were run for 1 μs each, and trajectories were saved every 25 ps.

### Metadynamics simulations

Well-tempered metadynamics simulations were performed using PLUMED [45] as interfaced with the AMBER18/AMBER20 simulation packages. The collective variables were chosen to be the extension and twist angles described earlier. Specifically, *extension* is defined to be the angle that connects the centers of mass of residues 376-391, 424-428, and 657-664. *Twist* is defined as the dihedral angle that connects the centers of mass of residues 376-391, 362-365, 424-428, and 657-664. Starting structures were taken from equilibrium MD simulations described after 500 ns of sampling. Gaussian hills were deposited every 500 integration timesteps with initial heights of 1.5 kJ/mol and a bias factor of 15. For the mon-omer-DNA complex system, we used multiple-walker metadynamics with twelve walkers exploring the collective variable space. Metadynamics simulations were considered converged when all the collective variable space had been sampled and free energy landscapes achieved asymptotic behavior with respect to number of hills included in the calculations.

### Post-simulation analysis

Principal components of motion were calculated using the ProDy [46] module in VMD [47]. Mutual information was calculated using the CARDS method [29] implemented in the Enspara package [48] with frames from both duplicate simulations, for a total of 2 μs of sampling saved in 80,000 frames for each nucleotide-bound state. Target site analysis was used to isolate the mutual information to locations of interest, as discussed in the results. VMD [47] and UCSF ChimeraX [49] were used for visualization. The Julia language [50] was used to create plots.

## Acknowledgements

We thank Drs. Marc Morais and Paul Jardine for valuable discussion pertaining to mechanistic implications of our simulations. We thank Joseph Magrino for critical reading of the manuscript. This work was partially supported by the National institutes of Health grant R01GM118817 (to G.A.). Computational resources were provided in part by the NSF XSEDE Program ACI-1053575. Anton 2 computer time was provided by the Pittsburgh Supercomputing Center (PSC) through Grant R01GM116961 from the National Institutes of Health. The Anton 2 machine at PSC was generously made available by D.E. Shaw Research. The authors declare no conflict of interest.

## Notes

### Competing Interest Statement

The authors have declared no competing interest.

