## Supplemental Info for "Molecular Dynamics of DNA Translocation by FtsK"

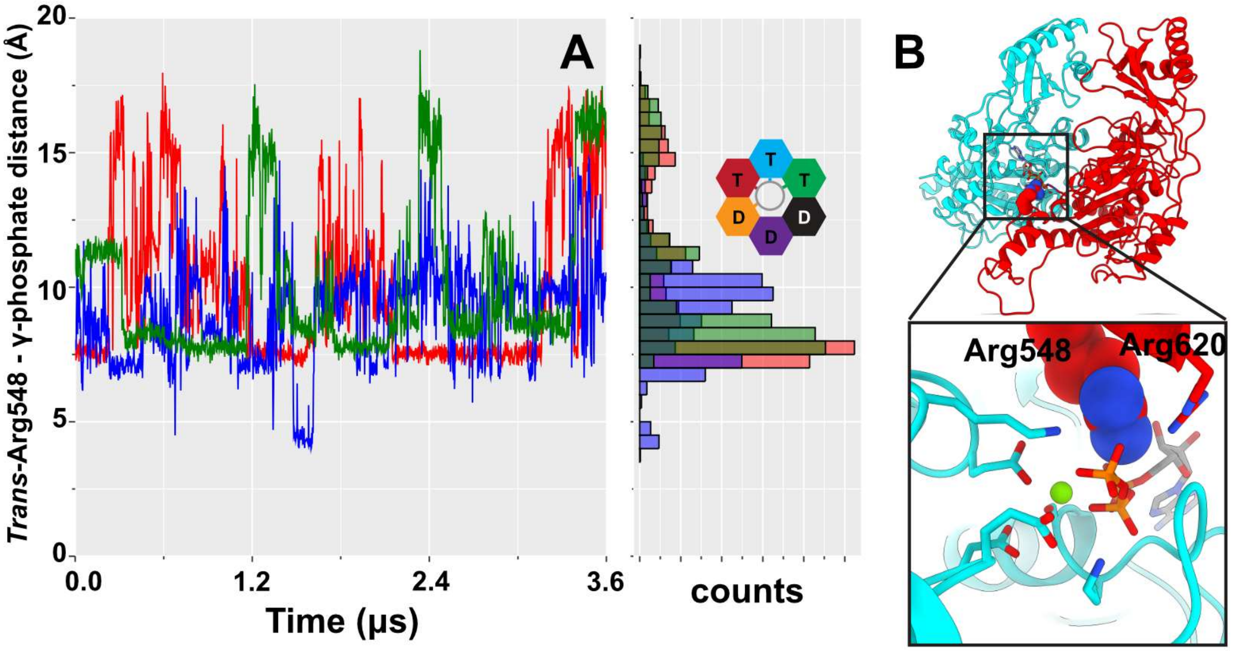


**Figure S1. Arg548 may also function as an additional arginine finger. (A)** Simulated distance between the guanidine carbon of Arg548 and the γ-phosphate of ATP in the neighboring subunit. Arg548 has the flexibility to briefly coordinate the γ-phosphate, suggesting that it could play a role in ATPase activity in addition to the proposed exchange role. Traces are colored according to the cis binding site. **(B)** A depiction of this interaction. The canonical arginine finger, Arg620, is also shown.

**Movie S1.** The first principal component of motion of the hexamer complex predicted from MD simulation. Subunit coloring is consistent with the coloring used in the manuscript (see figures 2, 3, 6, S1). DNA is shown static for reference. The trajectory was aligned about the β-ring. The first mode of motion shows that subunit motions are sympathetic.

**Movie S2.** The first principal component of motion of the hexamer complex predicted from MD simulation. For clarity, only the most extended ATP-bound subunit (green), and the most compact ADP-bound subunit (orange) are shown. DNA is shown static for reference. Subunits are aligned about the β-ring and show sympathetic motion in the α-ring. The large motion of the "exchange residue" tail can be seen in-plane in the green subunit; this motion is proposed to promote *trans*-activate nucleotide exchange.

**Movie S3.** The first principal component of motion of the hexamer complex predicted from MD simulation. Two subunits are shown, the ADP-bound subunit at the top of the β-ring (orange in the main text), and the neighboring ADP-bound subunit (purple in the main text). ADP is shown as a surface for reference. The trajectory was aligned about the α-ring. Subunits are colored according to root-mean-square fluctuations (red less flexible and blue more flexible) in order to highlight the flexibility of the exterior tail where Arg548 is located.
